# Amino Sugars Modify Antagonistic Interactions between Commensal Oral Streptococci and *Streptococcus mutans*

**DOI:** 10.1101/551168

**Authors:** Lulu Chen, Brinta Chakraborty, Jing Zou, Robert A. Burne, Lin Zeng

**Author notes:** Communicating author: 1395 Center Drive, D5-27, Gainesville, Florida 32611.

## Abstract

N-acetylglucosamine (GlcNAc) and glucosamine (GlcN) enhance the competitiveness of the laboratory strain DL1 of *Streptococcus gordonii* against the caries pathogen *Streptococcus mutans*. Here we examine how amino sugars affect the interaction of five low-passage clinical isolates of abundant commensal streptococci with *S. mutans* utilizing a dual-species biofilm model. Compared to glucose, growth on GlcN or GlcNAc significantly reduced the viability of *S. mutans* in co-cultures with most commensals, shifting the proportions of species. Consistent with these results, production of H_2_O_2_ was increased in most commensals when growing on amino sugars, and inhibition of *S. mutans* by *Streptococcus cristatus, Streptococcus oralis,* or *S. gordonii* was enhanced by amino sugars on agar plates. All commensals except *S. oralis* had higher arginine deiminase activities when grown on GlcN, and in some cases GlcNAc. In *ex vivo* biofilms formed using pooled cell-containing saliva (CCS), the proportions of *S. mutans* were drastically diminished when GlcNAc was the primary carbohydrate. Increased production of H_2_O_2_ could account in large part for the inhibitory effects of CCS biofilms. Surprisingly, amino sugars appeared to improve mutacin production by *S. mutans* on agar plates, suggesting that the commensals have mechanisms to actively subvert antagonism by *S. mutans* in co-cultures. Collectively, these findings demonstrate that amino sugars can enhance the beneficial properties of low-passage commensal oral streptococci and highlight their potential for moderating the cariogenicity of oral biofilms.

**SIGNIFICANCE:** Dental caries is driven by dysbiosis of oral biofilms in which dominance by acid-producing and acid-tolerant bacteria results in loss of tooth mineral. Our previous work demonstrated the beneficial effects of amino sugars, GlcNAc and GlcN, in promoting the antagonistic properties of a health-associated oral bacterium, *Streptococcus gordonii,* in competition with the major caries pathogen *Streptococcus mutans.* Here we investigated 5 low-passage clinical isolates of the most common streptococcal species to establish how amino sugars may influence the ecology and virulence of oral biofilms. Using multiple *in vitro* models, including a human saliva-derived microcosm biofilm, experiments showed significant enhancement by at least one amino sugar in the ability of most of these bacteria to suppress the caries pathogen. Therefore, our findings demonstrated the mechanism of action by which amino sugars may affect human oral biofilms to promote health.

## INTRODUCTION

Hundreds of bacterial taxa colonize the surfaces of the oral cavity in three-dimensional matrices termed biofilms (1), the composition of which is driven by adhesive interactions, a variety of synergistic and antagonistic interactions between the microorganisms, and host genetics and behaviors (2). Collectively, these factors determine the pathogenic potential of oral biofilms (3, 4). Numerous studies now support that the development of the most common oral infectious diseases, dental caries and periodontal diseases, is characterized by a change in the composition and biochemical activities of the microbiota in oral biofilms; from one that is rich in health-associated commensals to biofilms that contain substantially increased proportions of opportunistic pathogens or pathobionts. While the factors that drive the formation of periodontopathic biofilms are not fully understood, dental caries results from the establishment of strongly acidogenic (acid-producing) and aciduric (acid-tolerant) biofilms that are selected for by repeated ingestion by the host of fermentable carbohydrates and the resultant recurrent acidification of the biofilms by the organic acids produced during sugar fermentation (5).

*Streptococcus mutans* is considered a major etiologic agent contributing to the initiation and the progression of dental caries (6). One primary virulence attribute of *S. mutans* is extreme acidification of the environment from the fermentation of an array of carbohydrates (7, 8). Another determining factor that enables *S. mutans* to become a successful cariogenic bacterium is its exceptional capacity to form biofilms on teeth, largely facilitated by its robust production of extracellular polymeric substances (EPS) catalyzed by secreted glucosyltransferases (Gtfs) and fructosyltransferase (Ftf) enzymes that generate diffusion-limiting exopolysaccharides (6), and the ability to produce substantial quantities of extracellular DNA (eDNA) (9). The metabolic activities and the matrix combine to create localized low-pH environments that are ideal for *S. mutans* or other aciduric species to thrive, while these environments suppress the growth of health-associated commensal organisms, which are acid-sensitive compared to cariogenic organisms (2). Furthermore, strains of *S mutans* produce multiple lantibiotic and/or non-lantibiotic bacteriocins, collectively referred to as mutacins, which can inhibit the growth of a variety of Gram-positive bacteria (10). While direct evidence from *in vivo* studies is lacking, it appears that mutacins may be critical for allowing *S. mutans* to establish, persist and compete with commensal and overtly beneficial bacteria, especially oral streptococci (11). The ComDE two-component system and its cognate signal, competence-stimulating peptide (CSP), comprise the primary quorum sensing regulatory circuit controlling bacteriocin gene activation, although multiple other factors affect the production of mutacins (12).

As the most abundant species in many dental biofilms, commensal oral streptococci deploy multiple antagonistic strategies against pathogens, creating conditions that are favorable to dental health. For example, in the presence of oxygen *Streptococcus sanguinis*, *Streptococcus gordonii* and other members of the Mitis group of streptococci produce substantial (mM) quantities of hydrogen peroxide (H_2_O_2_), which has a potent inhibitory effect on the growth and physiology of *S. mutans* (4). Likewise, many of the Mitis group of streptococci express the arginine deiminase (AD) pathway, which moderates acidification of oral biofilms by releasing ammonia and carbon dioxide, while concurrently providing bioenergetic benefits to the producing organisms (13). All *S. mutans* lack the AD system. In addition, certain oral streptococci can interfere with intercellular communication systems in a way that reduces the production of mutacins by *S. mutans* and subverts the expression of other key virulence-related phenotypes, including genetic competence. For example, a novel commensal, designated *Streptococcus* A12, isolated from a caries-free human interferes with the CSP-ComDE signaling system required for mutacin production and the XIP (*comX*-inducing peptide) signaling pathway in *S. mutans* that directly regulates development of genetic competence (14).

Dietary carbohydrates are essential determinants of the cariogenic potential of dental biofilms (15, 16). Interestingly, analysis of the microbial composition of the fossil record and ancient calcified dental plaque indicates that dental caries and cariogenic bacteria, respectively, were not common until humans transitioned from a hunter-gatherer lifestyle to diets richer in natural and refined carbohydrates (17). A western diet, rich in carbohydrates, fuels caries development by greatly increasing the amount and frequency of acid production by oral biofilms. Data are now emerging in support of the notion that certain carbohydrates and end products may alter biofilm ecology by influencing the antagonistic relationships between health-associated commensals and caries pathogens. For example, glucose, a preferred carbohydrate source, at high concentrations represses H_2_O_2_ production by *S. gordonii* and reduce its competitiveness against *S. mutans* (4). *S. gordonii* significantly reduces H_2_O_2_ production under acidic conditions by diverting carbon flux toward lactic acid production and away from peroxidogenic enzymes, e.g. pyruvate oxidase (18). Similarly, AD expression in many streptococci is sensitive to carbon catabolite repression (CCR) (19). Conversely, certain amino sugars have been shown to enhance the competitiveness of *S. gordonii* against *S. mutans* via multiple mechanisms (20).

Amino sugars, including glucosamine (GlcN) and N-acetylglucosamine (GlcNAc), are extensively distributed in nature (21), serving as the building blocks for chitin, glycoproteins, bacterial peptidoglycans, and other abundant macromolecules (22, 23). Significant quantities of amino sugars and other carbohydrates are also present in human saliva (24–26), and can serve as carbon, nitrogen and energy sources for many members of the oral microbiota, including oral streptococci. In recent decades, GlcN and GlcNAc have become one of the most widely consumed dietary supplements in the U.S. and worldwide. Since amino sugars can be rapidly degraded by many oral streptococci and their catabolism leads to release of significant quantities of ammonia, which can raise cytoplasmic and environmental pH, metabolism of these sugars by the oral flora is expected to significantly influence oral biofilm composition and behaviors. Previous work by our group has indicated that metabolism of amino sugars benefits *S. mutans* by improving its acid tolerance (21). However, when compared to *S. mutans*, many commensal streptococci have superior abilities to utilize GlcN and especially GlcNAc, which is more common in nature. Also, unlike for *S. mutans,* GlcNAc can elicit catabolite repression in certain commensal oral streptococci as effectively as glucose (27). As a model commensal, *S. gordonii* DL1 outcompeted *S. mutans* in both planktonic and biofilm models, and was particularly dominant when amino sugars were provided (20). Preliminary genetic analysis also suggested that the cause of the dominance by *S. gordonii* included more efficient induction of the genes (*nag*) for the catabolic enzymes and increased expression of the AD pathway (20). However, questions remain as to whether similar influences of amino sugars exist on the other species of abundant commensal streptococci, as well as on complex oral biofilm communities. And, if so, what are the mechanisms and taxa that are having the most profound impact on the beneficial or detrimental behaviors of biofilm communities?

Here we selected a group of bacteria that were recently isolated from supragingival dental plaque of caries-free children and analyzed their interactions with *S. mutans* in the presence and absence of amino sugars. Further, using an *ex vivo* biofilm model derived from pooled whole-saliva, we examined the effects of amino sugars on the persistence and competitive fitness of *S. mutans* in a microcosm community. The results clearly demonstrate that GlcN and GlcNAc have the potential to modulate the cariogenic potential of dental biofilms, but the effects are complex and highly dependent on the types and proportions of health-associated oral bacteria that are present in the biofilms.

## RESULTS

### Utilization of amino sugars for growth by low-passage isolates of commensals

Previous studies by our group demonstrated the ability of a variety of oral streptococci, including laboratory reference strains, to utilize amino sugars for growth (20). To achieve a more comprehensive understanding of metabolism of amino sugars by dental biofilms, we obtained five clinical isolates derived from the supragingival plaque of caries-free subjects, each representing an abundant species of oral streptococci based on their 16S rRNA sequence: *Streptococcus sanguinis* BCC04, *S. gordonii* BCC09, *Streptococcus cristatus* BCC13, *Streptococcus oralis* BCC02, and *Streptococcus intermedius* BCC01. Phylogenomic analysis of the genome sequences of these strains has confirmed the assignment of species (28). As part of an ongoing study dissecting probiotic mechanisms of beneficial oral bacteria, these strains were assessed for traits that have been associated with dental health, including expression of the arginine deiminase pathway and antagonism of growth of *S. mutans* (Table 1).

**TABLE 1.**
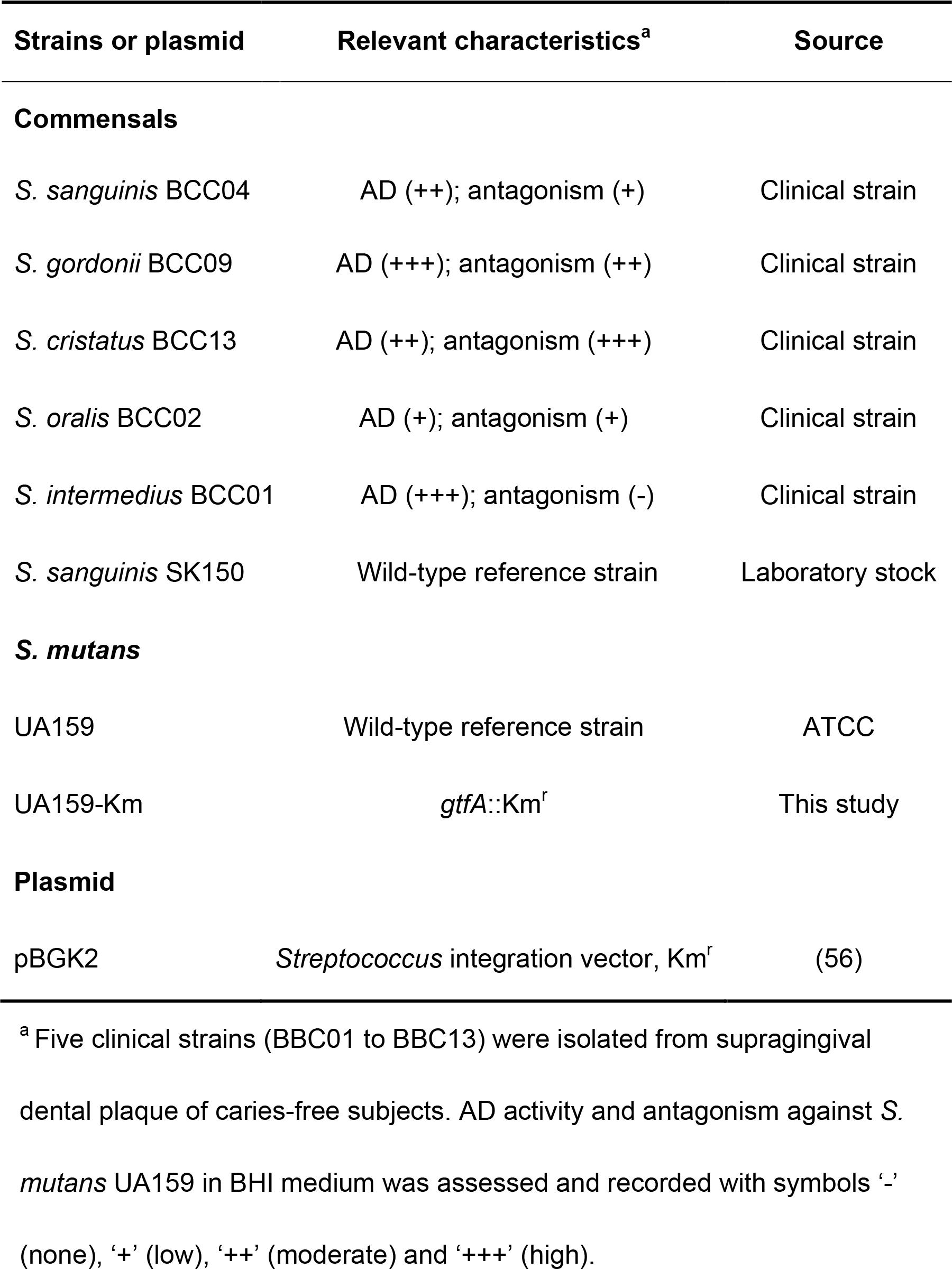
Strains and plasmids used in this study.

| Strains or plasmid | Relevant characteristics<sup>a</sup> | Source |
| --- | --- | --- |
| **Commensals** |  |  |
| *S. sanguinis* BCC04 | AD (++); antagonism (+) | Clinical strain |
| *S. gordonii* BCC09 | AD (+++); antagonism (++) | Clinical strain |
| *S. cristatus* BCC13 | AD (++); antagonism (+++) | Clinical strain |
| *S. oralis* BCC02 | AD (+); antagonism (+) | Clinical strain |
| *S. intermedius* BCC01 | AD (+++); antagonism (-) | Clinical strain |
| *S. sanguinis* SK150 | Wild-type reference strain | Laboratory stock |
| ***S. mutans*** |  |  |
| UA159 | Wild-type reference strain | ATCC |
| UA159-Km | *gtfA*::Km<sup>r</sup> | This study |
| **Plasmid** |  |  |
| pBGK2 | *Streptococcus* integration vector, Km<sup>r</sup> | (56) |

<sup>a</sup> Five clinical strains (BBC01 to BBC13) were isolated from supragingival dental plaque of caries-free subjects. AD activity and antagonism against *S. mutans* UA159 in BHI medium was assessed and recorded with symbols ‘-’ (none), ‘+’ (low), ‘++’ (moderate) and ‘+++’ (high).

First, to assess their capacity to utilize amino sugars, the five isolates were cultured to exponential phase in BHI broth, then diluted into FMC constituted with 20 mM glucose (Glc), GlcN, or GlcNAc. As shown in Fig. 1, *S. sanguinis* BCC04, *S. gordonii* BCC09 and *S. cristatus* BCC13 grew similarly well in all three carbohydrates, with only a modestly faster growth on Glc than on GlcNAc. By comparison, *S. oralis* BCC02 and especially *S. intermedius* BCC01 displayed significantly slower growth rates on GlcNAc, along with slightly longer lag phases and lower yields, compared to Glc or GlcN. Nevertheless, compared with the growth phenotype of *S. mutans* on GlcNAc (20), the commensal isolates showed no apparent defect in catabolizing amino sugars under these test conditions.

**FIGURE 1.**
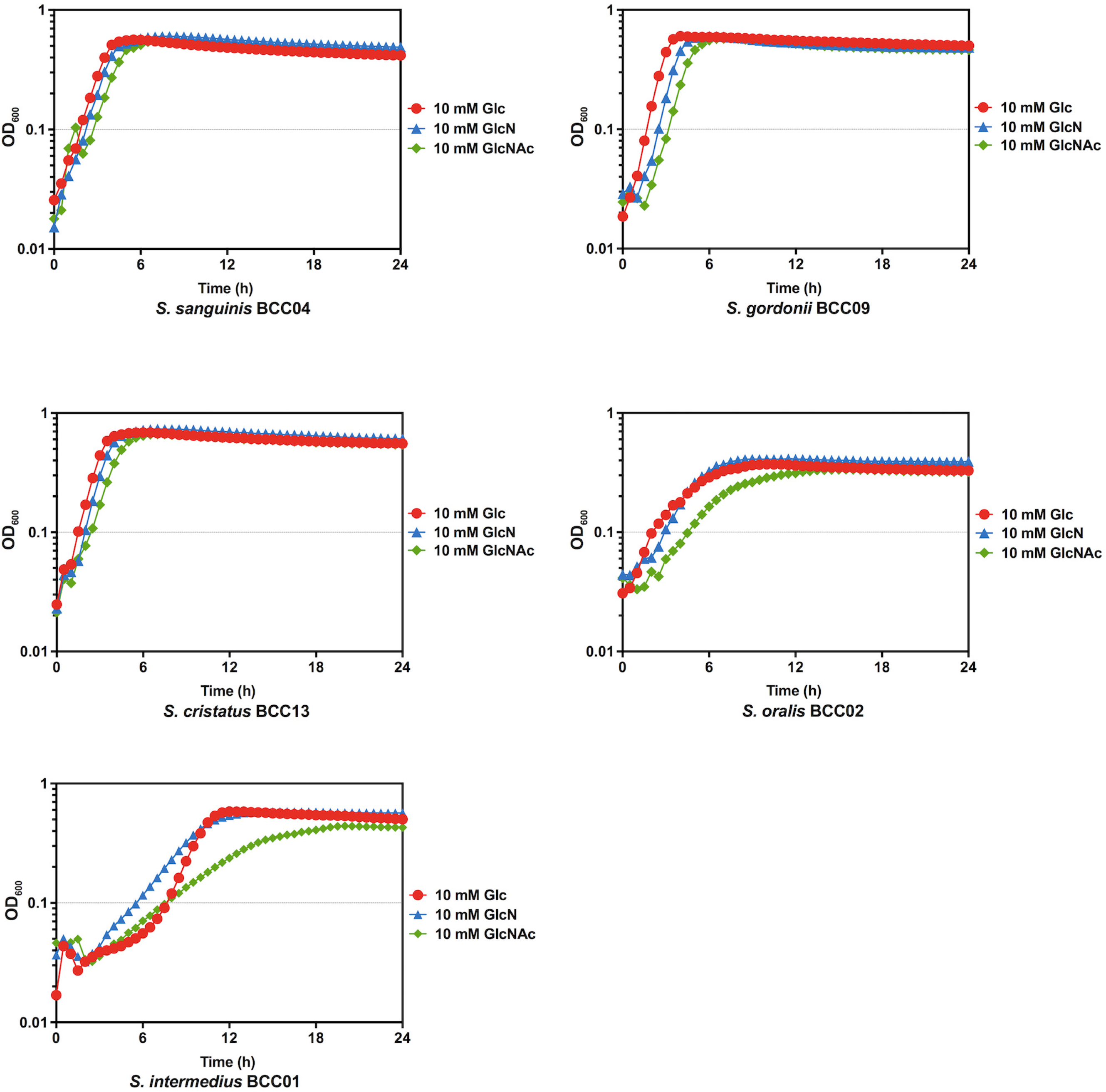
Growth curves. Commensal strains were cultured in BHI till the optical density at 600 nm (OD_600_) reached 0.5. Cultures were then diluted 1:50 into FMC medium supplemented with 10 mM Glc (red), GlcN (blue), or GlcNAc (green). The OD_600_ was monitored using a Bioscreen C with readings taken every 30 min.

### Amino sugars influence competition between *S. mutans* and commensals in biofilms

To investigate whether amino sugars can influence the interactions between the selected commensals and *S. mutans*, a dual-species biofilm model (20) was established. Bacterial cultures in exponential phase were prepared in BHI, and each commensal isolate, alone or in a 1:1 ratio with *S. mutan*s UA159-Km (29) was used to inoculate BMGS to form biofilms on glass surface.

*S. sanguinis* BCC04 formed mature biofilms with extensive structures, whereas the four other commensals produced only sparse microcolonies under these conditions, regardless of the primary carbohydrate source (Fig. S1). In contrast, *S. mutans* UA159 alone formed relatively thick biofilms, albeit consisting of significant proportions of dead cells (Fig. 2A). Further, *S. mutans* formed comparable amounts of biofilms on three different sugars, as indicated by both microscopy and the results of CFU enumeration (Table S1). We posit that the four commensals that did not form robust biofilms could not do so, at least in part, because they lack the ability to efficiently convert sucrose into water-insoluble glucans, which are known to enhance adherence to glass and biofilm maturation.

**FIGURE 2.**
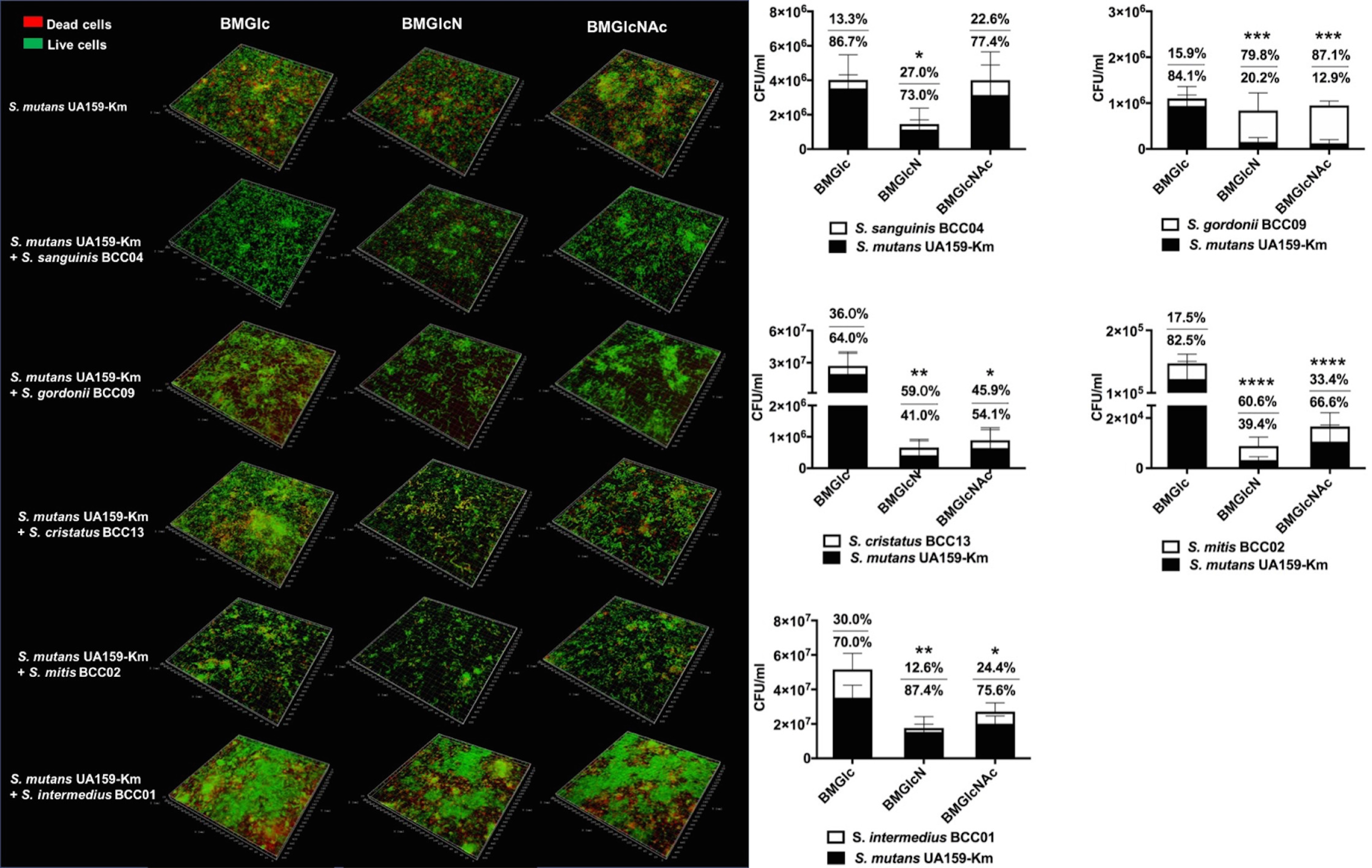
Dual-species biofilms formed using clinical commensals and *S. mutans*. Each commensal was mixed with UA159-Km in equal proportions and inoculated into BM supplemented with 18 mM Glc and 2 mM sucrose (BMGS) to form biofilms overnight on glass coverslips. BMGS was then replaced by BM supplemented with 20 mM Glc, GlcN, or GlcNAc and incubated for another 24 h before (A) visualization by confocal laser scanning microscopy (CLSM) after LIVE/DEAD staining and (B) CFU quantification of both species. Biofilms formed by UA159-Km alone were included for comparison. Values show the percentages of the populations constituted by commensal (top) and *S. mutans* (bottom) respectively. Asterisks indicate statistically significant differences in the viable counts of *S. mutans* compared to that on BMGlc, * *p* <0.05, ** *p* <0.01, *** *p* <0.001, **** *p* <0.001.

Even though different biofilm phenotypes were detected when *S. mutans* was combined with each of these commensals, some similar patterns were uncovered. Nearly all dual-species biofilms displayed the most extensive structures with Glc as the primary carbohydrate source, especially biofilms composed of *S. intermedius* BCC01 or *S. cristatus* BCC13 and *S. mutans* (Fig. 2A). In contrast, notably thinner biofilms were observed when the dual-species biofilms were formed using media formulated with GlcN or GlcNAc.

Consistent with their overall appearance, total CFU in these biofilms were strongly influenced by the carbohydrate source. Significantly lower CFU were obtained in dual-species biofilms composed of *S. cristatus* BCC13, *S. oralis* BCC02 or *S. intermedius* BCC01, together with *S. mutans,* when amino sugars were the supporting carbohydrate source. Co-cultures of *S. mutans* with *S. sanguinis* BCC04 produced significantly fewer total CFU on GlcN compared to Glc, however the results with Glc and GlcNAc were similar for BCC04.

The antibiotic marker engineered into strain UA159-Km allowed us to differentiate the two populations of bacterial cells in each system. In nearly all cases, when contrasted with Glc, one or both amino sugars yielded lower CFUs of UA159-Km in the biofilms, with significant reductions in the proportions of total CFU comprised of UA159-Km, except for *S. intermedius* (Fig. 2B). For example, amino sugars allowed *S. gordonii* BCC09 to become the dominant species in the dual-species biofilms (80% or higher), whereas the opposite was true with Glc. As an exception, co-cultures of *S. mutans* with *S. sanguinis* BCC04 showed no significant change in *S. mutans* cell counts on GlcNAc, relative to Glc.

### Growth on amino sugars increases H_2_O_2_ production by peroxidogenic commensals

It is believed that the H_2_O_2_ produced by commensal streptococci is a potent inhibitor of *S. mutans* and other oral pathobionts *in vivo* (30, 31). To begin dissecting the mechanisms by which amino sugars could improve the survival of commensals in co-cultures with *S. mutans*, the amount of H_2_O_2_ produced in liquid cultures by each commensal was quantified (Fig. 3).

**FIGURE 3.**
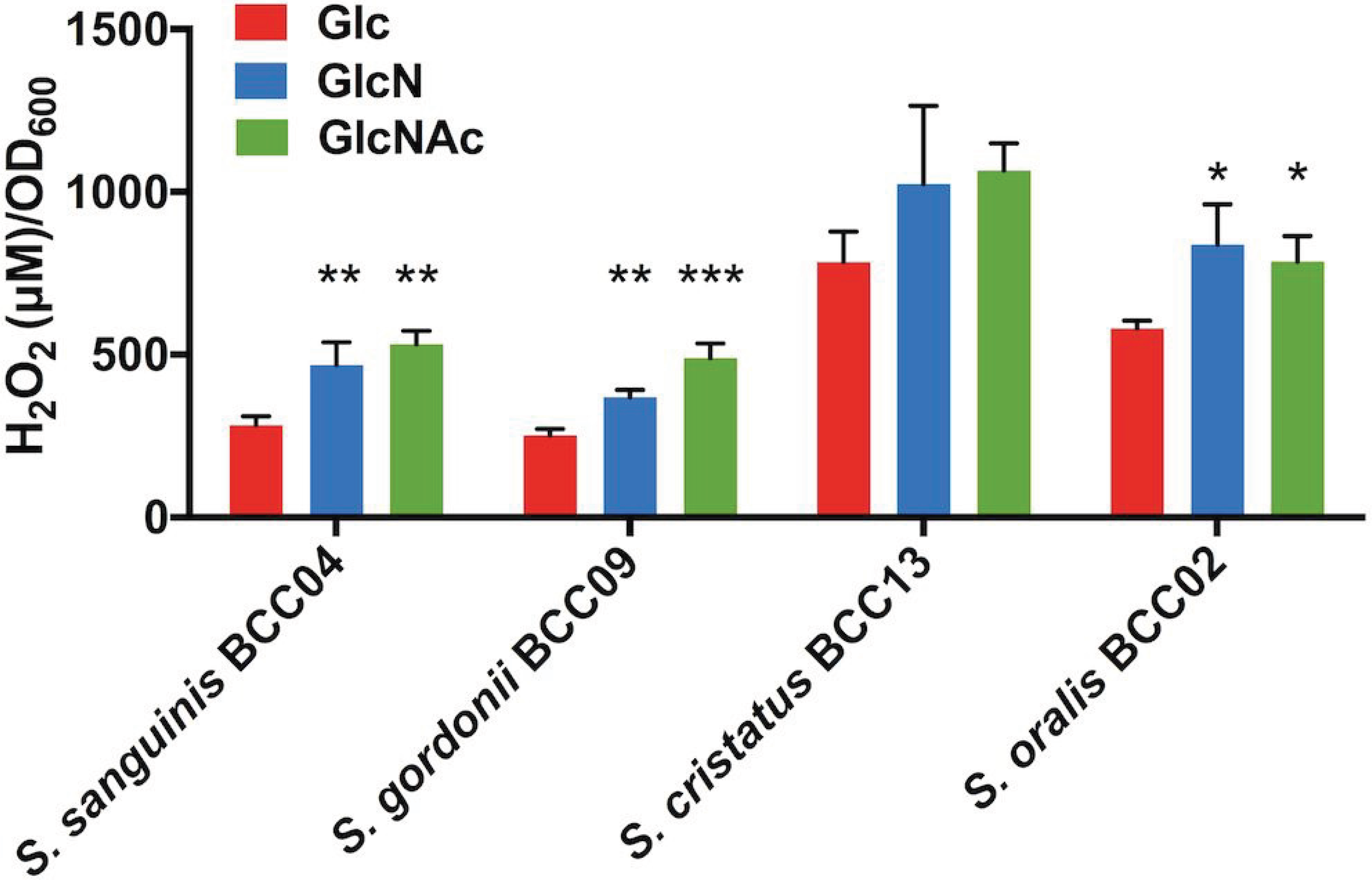
H_2_O_2_ production. Cells were cultivated to early exponential phase in TY supplemented with 20 mM Glc, GlcN or GlcNAc, exposed to vigorous aeration and the concentrations of H_2_O_2_ in the supernates were determined as described in the methods section. Asterisks indicate statistically significant differences in H_2_O_2_ levels relative to those on glucose, * *p* <0.05, ** *p* <0.01, *** *p* <0.001, **** *p* <0.001.

*S. gordonii* BCC09 produced significantly more H_2_O_2_ when growing in TY medium supplemented with amino sugars, especially GlcNAc, compared to cells growing on glucose (*P* <0.01). This finding is consistent with a previous report of a laboratory strain of *S. gordonii* (DL1) during its interactions with *S. mutans* (20). For *S. cristatus* BCC13 and *S. oralis* BCC02, relatively high concentrations of H_2_O_2_ were generated in all culture conditions, a result that is consistent with their outstanding ability to inhibit *S. mutans* in the biofilm model (Fig. 2). Importantly, more H_2_O_2_ was generated by BCC13 and BCC02 in the presence of either amino sugar, although some of these differences were not statistically significant. *S. intermedius* BCC01, on the other hand, was unable to produce detectable levels of H_2_O_2_ (data not shown), consistent with what has been described for some other *S. intermedius* strains (32). Interestingly, *S. sanguinis* strain BCC04 also produced more H_2_O_2_ when growing on either amino sugar, although it was only GlcN that enhanced its competitiveness in the biofilm model (Fig. 2B).

### Competition between *S. mutans* and clinical commensals on agar

Environmental factors such as oxygen tension and carbohydrate availability are known to not only affect the production of H_2_O_2_ by peroxidogenic species, such as *S. gordonii* (33), but also the production of mutacins (bacteriocins) by *S. mutans* (34). We performed a series of pair-wise competition assays to assess the effects of carbohydrate source and availability of oxygen on competition between the commensals and *S. mutans*. *S. mutans* UA159 and commensals were inoculated onto TY agar plates supplemented with 20 mM Glc, GlcN, or GlcNAc in three different ways: 1) *S. mutans* was spotted and incubated for 24 h before the commensal was spotted nearby, and plates were incubated an additional 24 h; 2) vice versa, the commensal was spotted first; and 3) both species were spotted simultaneously and incubated for 24 h.

As shown in Fig. 4, in an aerobic environment, *S. mutans* generally inhibited the growth of commensal bacteria when it was spotted first, irrespective of the carbohydrate source. However, when commensals were inoculated first, differential effects of the carbohydrate source on competition of commensals with *S. mutans* were evident. For example, *S. cristatus* BCC13 and *S. oralis* BCC02 showed significantly stronger inhibition of *S. mutans* on plates containing GlcN than Glc. For *S. gordonii*, the enhanced antagonism of *S. mutans* was particularly significant on GlcNAc-containing plates. In contrast, *S. sanguinis* BCC04 showed a slight, positive response to GlcNAc and *S. intermedius* BCC01 showed no inhibition of *S. mutans,* regardless of the carbohydrate source. Finally, when the commensals and *S. mutans* were spotted at the same time, *S. gordonii, S. cristatus* and *S. oralis* showed reduced evidence of antagonism, and only a modest effect of carbohydrate source was observed. Together, these results suggested that at least one amino sugar favored three commensals in antagonizing *S. mutans* on agar plates if the commensals were allowed to become established prior to plating of *S. mutans*.

**FIGURE 4.**
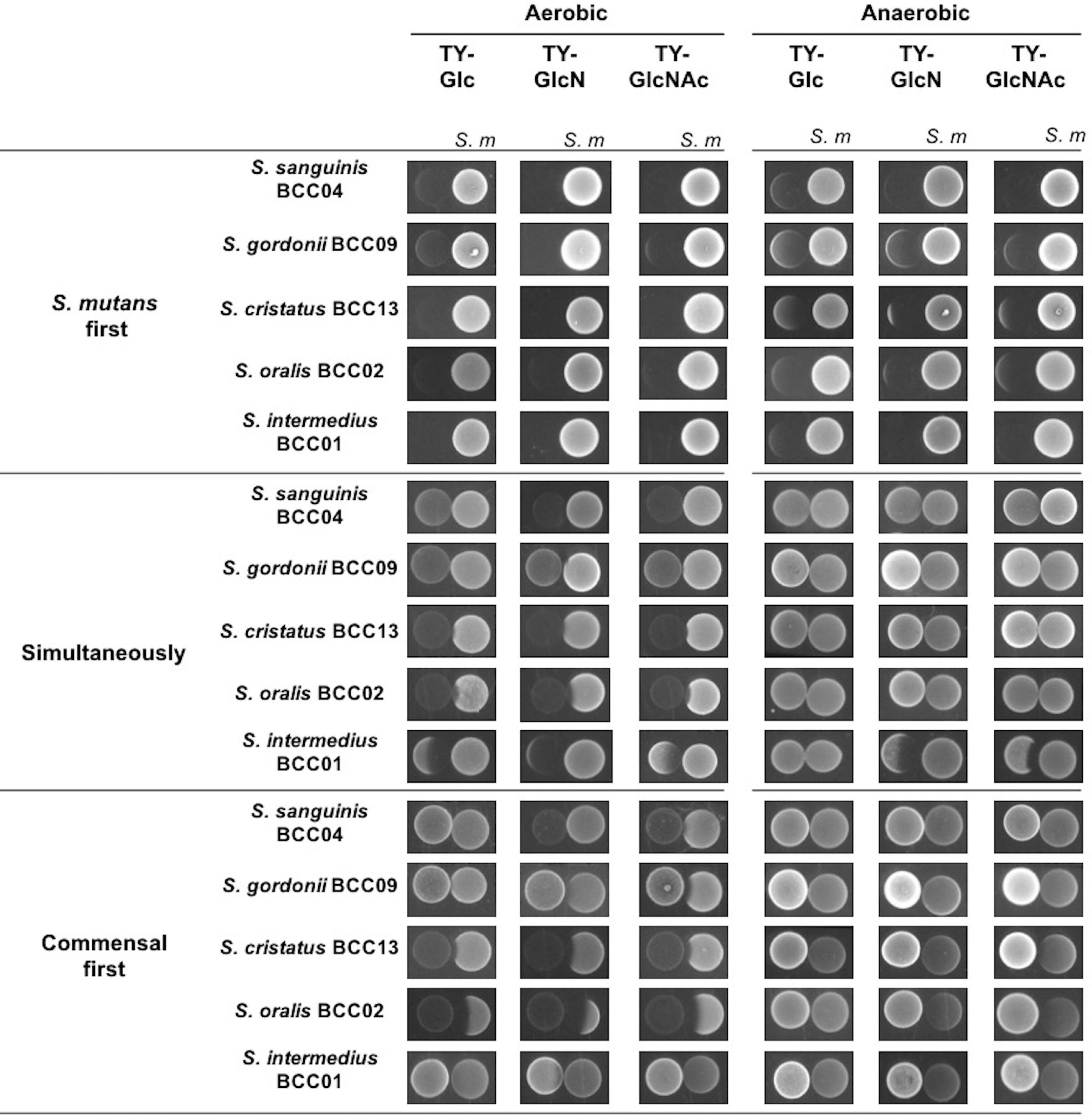
Competition assays between *S. mutans* and clinical commensals. Commensals and *S. mutans* UA159 were spotted on TY agar plates containing 20 mM Glc, GlcN, or GlcNAc, and incubated under aerobic or anaerobic conditions. The competitions were initiated with colonization by UA159 (*S. m* first), or commensals (Commensal first) alone for 24 h then followed by the other, or by both (Simultaneously).

To assess to what degree the presence of oxygen, and therefore the ability for the commensals to produce H_2_O_2_, contribute to these antagonistic activities, the same assays were conducted under anaerobic conditions. The results (Fig. 4) showed that most clinical commensals lost their ability to inhibit *S. mutans* regardless of the sequence of inoculation or carbohydrate source. Notably, when *S. mutans* was spotted first, it showed improved inhibition of all commensals when amino sugars were used in the plates, as compared to plates with glucose as the primary carbohydrate source. Thus, under more anaerobic conditions and/or in a reduced redox environment, the presence of amino sugars could potentially favor competition of *S. mutans* against beneficial commensals.

### Arginine deiminase (AD) activities by commensal isolates

In *S. gordonii* DL1, AD expression is repressed by glucose (19), but not by growth on amino sugars (20). To assess the effect of amino sugars on AD expression by the commensals in this study, the isolates were grown in TY medium containing 20 mM Glc, GlcN, GlcNAc or galactose, each supplemented with 10 mM arginine. The pH of the supernatant fluid was recorded after the cultures were harvested in exponential phase (Fig. S2). Galactose was used as a positive control as it does not significantly repress AD expression (35). For *S. sanguinis* BCC04, *S. cristatus* BCC13, and most notably *S. gordonii* BCC09 and *S. intermedius* BCC01, addition of GlcN resulted in enhanced AD activity, compared to cells growing on Glc (Fig. 5). AD activity was modestly higher on GlcNAc, albeit the differences were not statistically significant, for *S. gordonii* BCC09. Notably, the pattern of culture pH values for each strain corresponded with AD activity. Specifically, galactose-grown cultures presented the highest final pH values compared with other sugars; and for *S. gordonii, S. cristatus* and *S. intermedius*, use of amino sugars also resulted in elevated pH, compared to glucose-grown cultures (Fig. S2). The lack of efficient induction of AD in the presence of GlcNAc relative to GlcN has been observed for *S. gordonii* DL1 (20), however the molecular basis for this difference is not defined. *S. oralis* BCC02 produced negligible amounts of AD activity under all conditions and its overall pH values were significantly lower than the other strains.

**FIGURE 5.**
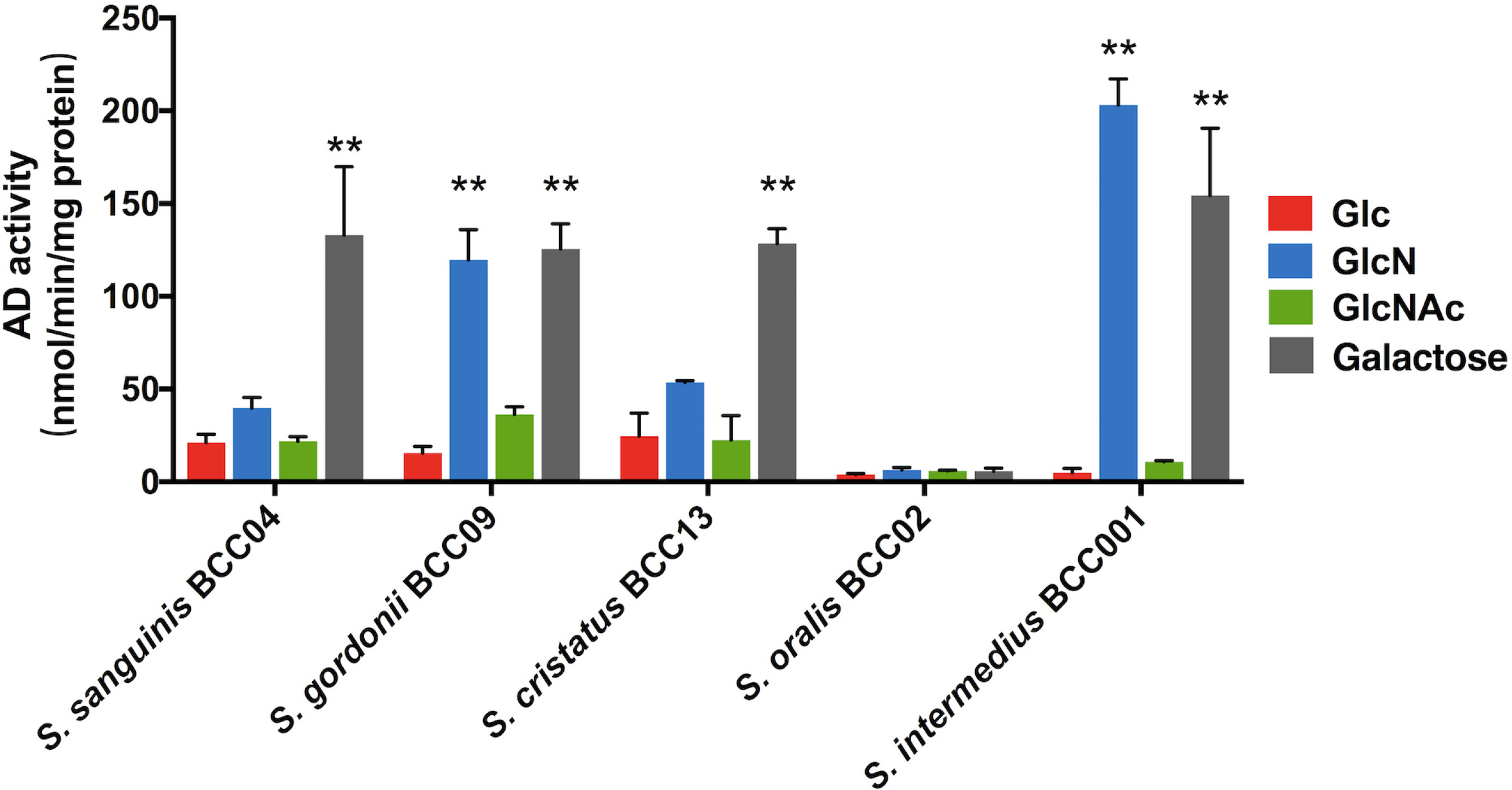
Arginine deiminase (AD) activity. For AD assays, commensals were grown to exponential phase in liquid TY supplemented with 20 mM Glc, GlcN, GlcNAc, or galactose, in addition to 10 mM arginine. Asterisks denote statistically significant difference compared to cultures prepared with glucose, * *p* <0.05, ** *p* <0.01, *** *p* <0.001, **** *p* <0.001.

### Bacteriocin production by *S. mutans* is affected by amino sugars

*S. mutans* secretes various antimicrobial peptides called mutacins that are capable of inhibiting the persistence of certain commensal streptococci (11, 36), an activity that can be countered by some commensals (14, 37). In order to investigate if production of mutacins is affected by amino sugars or the presence of the five organisms tested here, a soft-agar overlay assay was performed using *S. sanguinis* SK150 as the indicator strain (14). *S. mutans* UA159 alone, or UA159 together with one commensal, were inoculated on agar plates containing Glc, GlcN or GlcNAc. After 1 day of incubation, soft agar containing *S. sanguinis* SK150 cells was overlaid and the incubation was continued for 24 h before measuring zones of inhibition. The soft agar used to prepare SK150 suspensions was formulated to contain the same carbohydrate that was in the agar plates, and SK150 grew equally well on all three sugars (Fig. S3). As shown in Fig. 6, when *S. mutans* was used alone, it produced generally larger inhibition zones in aerobic conditions versus anaerobic incubation, when tested on the same carbohydrates. Interestingly, the largest zones of inhibition occurred on GlcN plates, followed by those on GlcNAc, which were slightly larger than those on Glc plates. These results were consistent with our previous study and may be associated with the effects of preferred carbohydrates, pH and oxygen on mutacin gene expression and activity (21).

**FIGURE 6.**
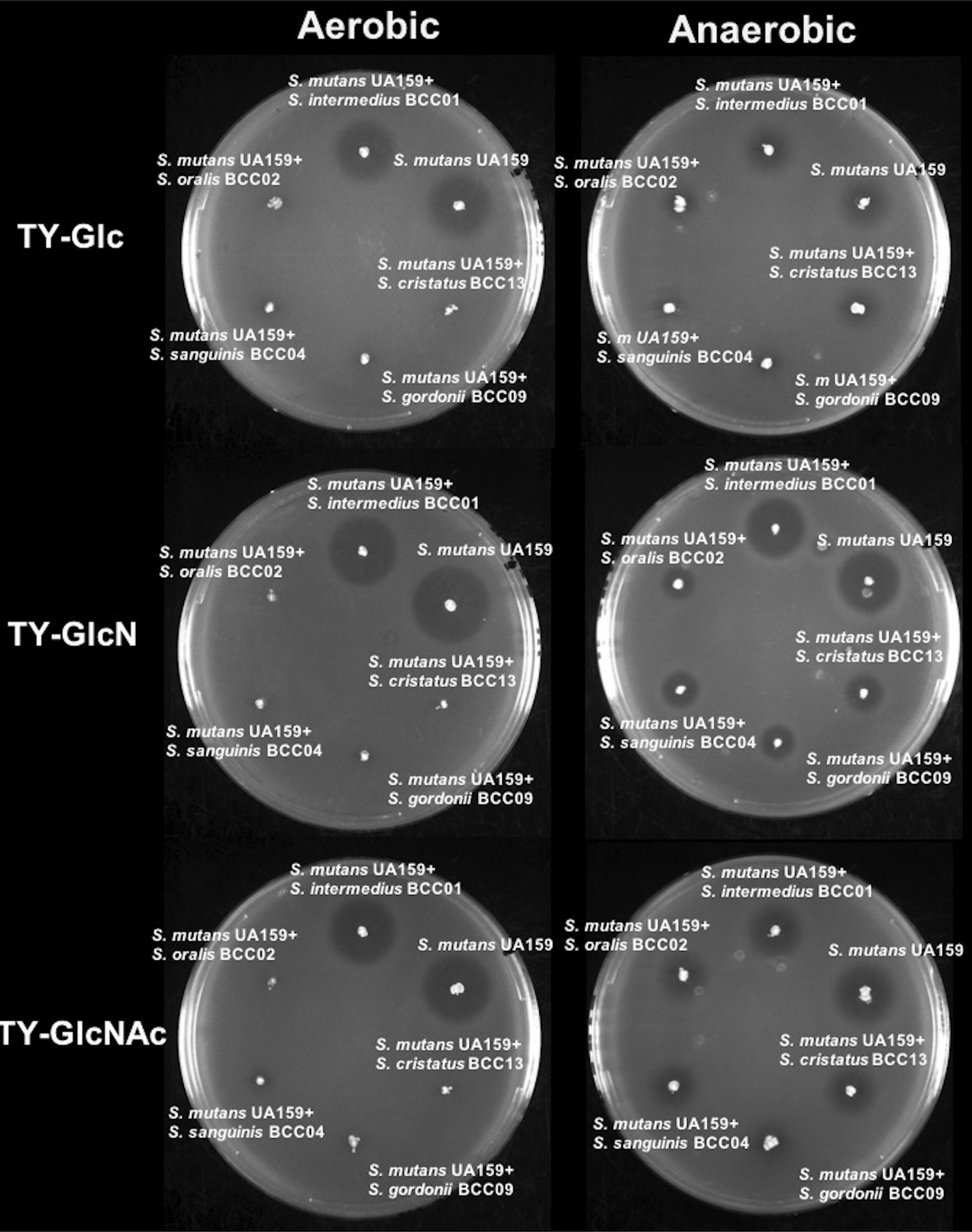
Plate-based bacteriocin assay. Washed cells of each clinical commensal were mixed with UA159 at a 1:1 ratio based on optical density and stabbed onto TY agar plates containing 20 mM Glc, GlcN, or GlcNAc. The plates were incubated overnight, overlaid with soft agar containing *S. sanguinis* SK150, and then incubated for another 24 h. The experiments were carried out in both aerobic and anaerobic environments.

Co-inoculation with commensals in aerobic conditions largely abolished the zone of inhibition of SK150 by *S. mutans* in all cases except for *S. intermedius*, regardless of the carbohydrate sources (Fig. 6). Conversely, incubation of the plates under anaerobic conditions significantly reduced, but did not completely remove, the effects of the four commensals on the reduction of the size of the zones of inhibition. These results were consistent with H_2_O_2_ being a dominant factor in the ability of the commensals to inhibit *S. mutans*. Different from the results of earlier assays (Figs. 2 and 4), among the four commensals that influenced the size of the zones of inhibition, similar levels of efficiency were observed under aerobic condition, regardless of the carbohydrate species. However under anaerobic conditions, these four effective strains showed the greatest impact on reducing the zone of inhibition when assayed on plates supplied with GlcN, followed by GlcNAc. Collectively, these results suggest that while H_2_O_2_ remains an effective mechanism for commensals to reduce the antagonistic capacity of *S. mutans,* additional factors are produced by particular commensals that are more dependent on carbohydrate source to affect production or effectiveness of mutacins by *S. mutans.*

### GlcNAc adversely affects persistence of *S. mutans* in an *ex vivo* biofilm model

*S. mutans* persists in a dynamic biofilm environment with hundreds of bacterial taxa, most of which have not been well-characterized (38–40). To better mimic the complex communities in the oral cavity, we employed an established *ex vivo* model system (41, 42) in which cell-containing whole saliva (CCS) samples were pooled from 4 donors and used to seed biofilms on glass surfaces. A genetically marked *S. mutans* strain (UA159-Km) was spiked into the biofilms and the effects of various sugars on persistence of *S. mutans* were monitored by CFU enumeration.

A schematic of the experimental design and various groups is shown in Fig. 7A. A biofilm medium (43) was used, but the carbohydrate content was modified such that Glc, GlcN or GlcNAc was present at 18 mM and 2 mM sucrose was present in all samples. In group *a*, *S. mutans* was co-inoculated with CCS in BMGS (Glc) for the first 24 h, then the medium was replaced with BMGlcNS (GlcN), BMGlcNAcS (GlcNAc), or BMGS for the next 24 h before plating. Compared with biofilms grown in BMGS only, the CFU of *S. mutans* in biofilms provided with GlcN was 3.6-fold greater, however CFU of *S. mutans* from GlcNAc-cultured biofilms were hardly detectable (Fig. 7B).

**FIGURE 7.**
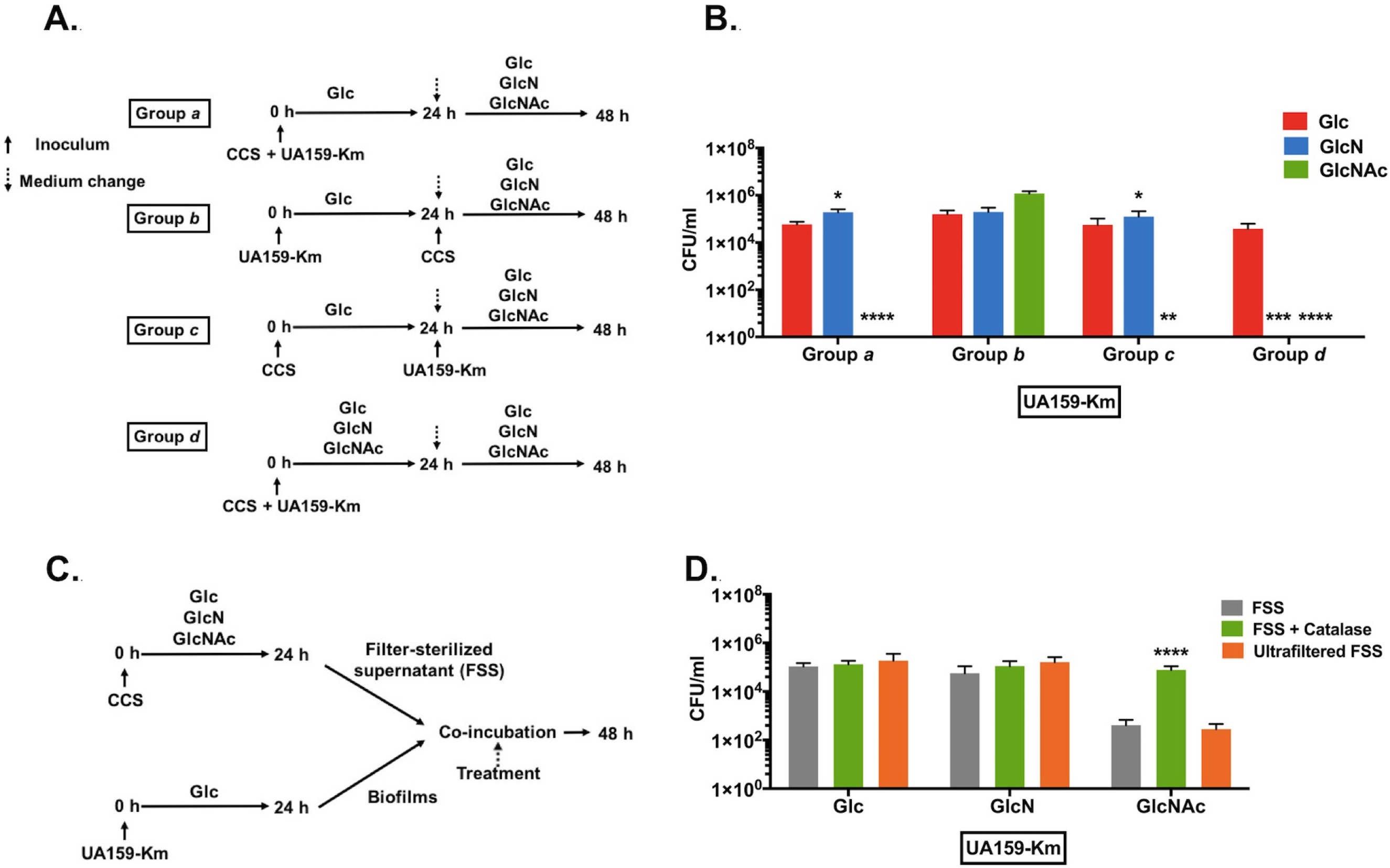
Amino sugars impact the survival of *S. mutans* in CCS-derived biofilms. (A, B) To initiate a biofilm, UA159-Km and cell-containing saliva (CCS) were used simultaneously (groups *a*, *d*), or 24 h apart (groups *b*, *c*) to inoculate BM medium supplemented with 2 mM sucrose and 18 mM various other carbohydrates. After 24 h of incubation, the biofilms were washed and resupplied with fresh media and additional bacteria as specified (see methods section for details). At the end of 2-day incubations, biofilms were processed to quantify the CFU of UA159-Km (B). Asterisks indicate statistically significant difference compared to cultures prepared with Glc, * *p* <0.05, ** *p* <0.01, *** *p* <0.001, **** *p* <0.001. (C, D) A UA159-Km biofilm was treated for 24 h with a filter-sterilized supernate (FSS) derived from biofilm cultures of CCS that were formed aerobically under different carbohydrate conditions, followed by CFU quantification of UA159-Km (D). The FSS was treated with catalase, passed through an ultrafiltration device (MWCO 10 kDa), or left untreated. Asterisks indicate statistically significant difference compared to untreated FSS, * *p* <0.05, ** *p* <0.01, *** *p* <0.001, **** *p* <0.001.

To determine whether the CCS inocula or *S. mutans* was able to first establish a biofilm, prior to inoculation with *S. mutans* or CCS, respectively, *S. mutans* (group *b*) or CCS (group *c*) was inoculated alone as the initial colonizer. Then CCS (*b*) or *S. mutans* (*c*), respectively, was introduced on the following day; carbohydrate sources were varied as for group *a*. When the initial colonizer was *S. mutans*, slightly higher CFU of *S. mutans* were recovered from cultures grown with GlcNAc, compared to Glc or GlcN (Fig. 7B, group *b*). In contrast, when the initial inoculum was CCS (group *c*), similar to what was seen in group *a*, very few *S. mutans* cells were detected in GlcNAc biofilms, whereas the CFU of *S. mutans* recovered from GlcN biofilms were comparable to Glc-cultured biofilms.

We then examined how the biofilms would mature if the amino sugars were present from the time of initial inoculation (group *d*). In this case, CCS and *S. mutans* were co-inoculated as group *a* and the cells were cultured in BM constituted with sucrose and either Glc, GlcN or GlcNAc for 48 h. The resultant cell counts of *S. mutans* were below the limit of detection in GlcNAc-grown biofilms. Of note, unlike the three other experiments, treatment with GlcN for 48 h also drastically reduced the CFU of *S. mutans* recovered from the biofilms (Fig. 7B). Clearly, for this particular biofilm model, addition of GlcNAc, and GlcN for longer periods of time, could result in profound changes in the ecology of the microbiome, significantly affecting the persistence of *S. mutans.*

Next, to determine if H_2_O_2_ or certain secreted factors were responsible for the diminished persistence of *S. mutans* associated with growth on amino sugars, UA159-Km alone was used to form biofilms on glass by incubating in BMGS for 24 h. 24-h CCS biofilms were similarly prepared using BMGS, BMGlcNS or BMGlcNAcS (Fig. 7C). The supernatant fluids of each CCS biofilm culture were harvested and filter-sterilized, then a portion of the sample (FSS) was treated with catalase to degrade H_2_O_2_ (FSS + Catalase) and another aliquot was passed through an ultrafiltration device to eliminate constituents with a molecular mass >10 kDa (Ultrafiltered FSS). Along with untreated control FSS, these different FSS samples were used to treat 1-day *S. mutans* biofilms for 24 h, followed by CFU enumeration of *S. mutans*. As shown in Fig. 7D, when FSS derived from Glc or GlcN cultures was added to the *S. mutans* biofilms, neither catalase treatment nor ultrafiltration had any notable impact on the survival of *S. mutans*. In stark contrast, for FSS prepared using GlcNAc-based cultures, catalase treatment significantly reduced the efficiency of FSS to inhibit the survival of *S. mutans,* whereas ultrafiltration had no effect.

Combining all the experimental settings, GlcNAc has been most effective in inhibiting *S. mutans* in this complex biofilm. Indeed, the highest amounts of H_2_O_2_ were measured in FSS from GlcNAc-based CCS biofilms, approximately 6 times that from GlcN-based biofilms (Fig. S4), reaffirming the role of H_2_O_2_ in amino sugar-dependent impacts on antagonism of *S. mutans* by oral commensals. In a control experiment, FSS derived from anaerobically formed CCS biofilms were used to treat the same 1-day *S. mutans* biofilm for 24 h. The resultant CFU counts of *S. mutans* showed no difference among three different carbohydrates (Fig. S5).

### DISCUSSION

The concept that a variety of abundant commensal bacteria, particularly oral streptococci, have overtly beneficial properties that include the ability to moderate biofilm acidification and to antagonize the growth of caries pathogens is central to the current understanding of the ecological underpinnings of dental caries. Antimicrobial compounds and fitness-promoting activities of the pathogen(s) and commensals are key components of the interactions that drive biofilm formation, foster community stability and lead to ecological changes during disease development. While dietary carbohydrates can drive the formation of caries, the majority of time the oral microbiome relies on host and bacterially derived nutrient sources, with amino sugars being an abundant and bioenergetically favorable source of carbon, nitrogen and energy. The goal of the present study was to assess the effects of two amino sugars, GlcN and GlcNAc, on some of the most abundant oral streptococcal species in the context of their overall fitness and antagonistic interactions with *S. mutans*. To that end, low-passage isolates from five representative abundant oral *Streptococcus* species were selected from recently obtained strains (28), rather than using standard lab strains that had been extensively passaged *in vitro*. The five isolates selected also displayed a spectrum of activities in phenotypes that have been deemed critical to antagonism and pH moderation. Using a number of established model systems, this study demonstrated the profound ability of amino sugars to influence interbacterial interactions that are central for shaping the ecological balance and virulence potential of dental biofilms. Specifically, amino sugars enhanced H_2_O_2_ production by peroxidogenic streptococci in liquid cultures and on agar plates (*S. oralis, S. cristatus* and *S. gordonii*). Moreover, commensal oral streptococci are generally considered less acid-tolerant than *S. mutans,* and growth on GlcN also resulted in significantly increased AD activity and culture pH in some of the commensals. Although not at all detrimental to *S. mutans,* production of ammonia via the AD system and from amino sugars can protect the organisms against acid stress. Thus, the combination of arginine and amino sugars may be quite beneficial to commensals in competition with *S. mutans* and other caries pathogens, which gain their advantage over the commensals by virtue of their ability to grow and metabolize at lower pH values than the commensals. Consistent with these trends, amino sugars largely favored the dominance and survival of commensals and diminished the persistence of *S. mutans* in dual-species and *ex vivo* consortium biofilm models.

A significant finding arising from this study is the distinct effects exerted by GlcN and GlcNAc on the selected commensals, and the significant differences in behaviors and outcomes noted among the models, which are designed to examine certain behaviors, albeit in different contexts. For example, addition of GlcN, as opposed to GlcNAc, enhanced AD activity by most clinical commensals in liquid culture (Fig. 5). Furthermore, GlcN was notably more effective at stimulating anti-*S. mutans* activities on plates by *S. cristatus* and *S. oralis* under aerobic conditions, apparently due to H_2_O_2_ production. The presence of either amino sugar in liquid cultures resulted in comparable enhancements in H_2_O_2_ production by almost all the clinical commensals. Although GlcN promoted the secretion of mutacins by *S. mutans* to a greater extent than did GlcNAc (Fig. 6), overall these results provide support for the idea that GlcN is the better amino sugar for enhancing the competitiveness of the chosen commensals.

However, in the saliva-derived *ex vivo* biofilm model, where a far more complex community exists to compete against *S. mutans*, the biofilm as a whole responded favorably to GlcNAc against colonization by the pathogen, whereas GlcN appeared to benefit commensals only if present for longer periods of time. Thus, it is apparent that the two amino sugars of interest can exert different impacts on the behaviors of both the commensals and *S. mutans.* It must be stressed that these *in vitro* models differ from one another in significant ways: they differ in oxygen tension, nutrient availability, colonization sequence, and biodiversity. In particular, saliva is a natural reservoir of indigenous bacteria found in oral cavity. Not only does saliva harbor a microbial community that closely matches that of dental plaque in a given individual (44), it contains factors that strongly influence the development and maintenance of the dental microbiome, including mucins, trace elements and host- and microbiota-derived enzymes (38). Future studies will focus on exploring the mechanisms responsible for the distinct effects of amino sugars in the context of oral biofilm complexity and host factors.

While studying a complex biological system such as the dental biofilm, a reductive approach comes with benefits and potential deficiencies. Genomic heterogeneity and functional overlap in the microbiota have proven widespread among the numerous streptococcal species colonizing the oral cavity (45). It is generally accepted that exchange of genetic material is a common occurrence among many members of the oral microbiome, especially among streptococci. In this study, we have noted that the low-passage isolate *S. sanguinis* BCC04 was partly responsive to amino sugars (GlcN only) in the dual-species biofilm model (Fig. 2) and inhibited mutacin production (Fig. 6) in a manner that is independent of oxygen. In particular, BCC04 showed no enhancement in its antagonism against *S. mutans* in the presence of amino sugars on plates (Fig. 4) and in the biofilms supported by GlcNAc, despite the fact that it produced significantly more H_2_O_2_ when supplied with either amino sugar. As a complement to these experiments, a commonly used lab strain *S. sanguinis* SK150 was tested in the same dual-species biofilm model. The results showed a differential response to carbohydrates, with both amino sugars yielding significantly lower viable counts of *S. mutans* (Fig. S6). These disparate phenotypes of *S. sanguinis* isolates are likely a result of intra-species genomic heterogeneity that affects their competition with *S. mutans*. In fact, strain BCC04 produces robust biofilms in the presence of sucrose (Fig. S1), whereas SK150 does not (data not shown). Recently, it was reported that another laboratory strain of *S. sanguinis,* SK36, showed a lack of response to carbohydrate availability in its ability to produce H_2_O_2_ (46), echoing a previous finding that *spxB* expression in *S. sanguinis* was unresponsive to carbohydrate source (47). Comparative and functional genomic analyses currently being performed on these and other recent isolates of *S. sanguinis* could shed more light on the cause of these distinct phenotypes. In a separate study that focused on the effects of arginine on the antagonistic interactions between oral commensals and *S. mutans*, it was shown that enormous diversity exists among commensal streptococci, both in their antagonistic capabilities and their susceptibility to *S. mutans* (48). Clearly, probing the basis for phenotypic variation among commensals and pathogens (49) remains an essential goal in efforts to improve caries risk assessment and intervention.

Another notable finding from this study is the superior capacity of *S. cristatus* BCC13 and *S. oralis* BCC02 to antagonize *S. mutans*; and both strains showed improved abilities to inhibit *S. mutans* when amino sugars were present (Fig. 2&4). While limited research has been conducted on the interactions between *S. cristatus* and *S. mutans,* it is established that *S. cristatus* is able to adversely affect the periodontal pathogen *Porphyromonas gingivalis* (Pg), both by repressing the expression of the primary fimbrial adhesin of Pg (50) and through production of H_2_O_2_ (51). As shown here, *S. cristatus* is clearly capable of disrupting the persistence of *S. mutans* by producing large amounts of H_2_O_2_, and potentially reducing the production of mutacins. Some recent reports have indicated that *S. cristatus* could be associated with severe early childhood caries, however more research is needed to corroborate these findings (52, 53). Similarly, *S. oralis* BCC02 appeared highly inhibitory to *S. mutans* in biofilms, at least partly due to being an efficient H_2_O_2_ producer. *S. oralis* has been reported to maintain homeostasis in oral biofilms, primarily by suppressing *S. mutans* overgrowth (54) and influencing biofilm formation by *S. mutans* (43). Earlier studies also demonstrated the remarkable ability of *S. oralis* to produce hydrogen peroxide (55), which is corroborated by the present study.

In conclusion, this study demonstrated the capacity of amino sugars in promoting endogenous commensal bacteria to antagonize a major dental caries pathogen, *S. mutans*. It is also clear that the effects of amino sugars on streptococci, both the pathogen and commensals alike, are quite complex. For example, while *S. mutans* alone has been shown to respond favorably to the presence of amino sugars by enhancing aciduricity (21) or the production of mutacins, co-cultivation with the commensal species under the same conditions has resulted in greater reductions in persistence of the pathogen compared to glucose. Aside from the enhanced production of H_2_O_2_ by the commensals, another possible explanation to this outcome is that commensals actively subvert the competitiveness of *S. mutans* when amino sugars are present in sufficient concentrations (14). While we expand our efforts to identify amino sugar-responsive, antagonistic commensals, availability of genomic information of these isolates (28) will greatly aid in our understanding of the basis for phenotypic behaviors affected by amino sugars. Based on this and other studies, amino sugars may have benefits when included in prebiotic or synbiotic strategies for caries management.

## MATERIALS AND METHODS

### Bacterial strains and culture conditions

The bacterial strains used in this study are listed in Table 1. A series of clinical isolates belonging to different *Streptococcus* species were selected from caries-free plaque samples and characterized by 16S rDNA sequencing. Five such isolates, *S. sanguinis* BCC04, *S. gordonii* BCC09, *Streptococcus cristatus* BCC13, *Streptococcus oralis* BCC02, and *Streptococcus intermedius* BCC01, were also screened for their abilities to express the arginine deiminase (AD) enzyme and to antagonize

*S. mutans*. All streptococcal strains, including *S. mutans* UA159, and a mutant derivative UA159-Km with a kanamycin resistance cassette that replaced the non-essential sucrose phosphorylase (*gtfA*) gene, by using plasmid pBGK2 (56). Transformants were selected on brain heart infusion (BHI) (Difco Laboratories, Detroit, MI) agar containing kanamycin (1 mg/ml) and maintained on BHI agar plates. Liquid BHI was used to prepare starter cultures for assays. TY (3% Tryptone, 0.5% yeast extract) medium was used to grow cells for measurements of H_2_O_2_ production and AD activity, and as the base for agar plates used in a competition assays and a bacteriocin production assay. TY medium, the chemically defined medium FMC (21) and a biofilm medium BM (43) were each formulated to contain various carbohydrates at specified concentrations. Bacterial planktonic or biofilm cultures in most experiments were incubated at 37°C in a 5% CO_2_, aerobic environment or in anaerobic jars (BD, Franklin Lakes, NJ) as noted.

### Planktonic growth assays

Commensals were cultured overnight in BHI and sub-cultured in BHI broth to mid-exponential phase (OD_600_ ≈ 0.5), then diluted 1:50 into modified FMC medium formulated with final concentrations of 10 mM glucose (Glc), glucosamine (GlcN) or N-Acetyl-D-glucosamine (GlcNAc) (Sigma Millipore, St. Louis, MO). Cultures were maintained at 37°C, and the optical density at 600 nm (OD_600_) was measured and recorded every 30 min using a Bioscreen C Lab system (Helsinki, Finland) for 24 h. To reduce exposure to air, approximately 60 μl of mineral oil was overlaid on each sample (57).

### Biofilm assays

Overnight bacterial cultures were diluted 1:20 into fresh BHI medium and sub-cultured to OD_600_ ≈ 0.5. Each culture of the commensal alone, or together with *S. mutans*-Km in a 1:1 ratio, was prepared by diluting (1:100) into a final volume of 400 μl of biofilm medium (BM) (43) that contained 18 mM glucose (Glc) and 2 mM sucrose (S). The cultures were kept in an Ibidi glass plate (Ibidi USA Inc., Madison, WI). After 24 h of incubation in aerobic conditions, the spent medium was removed, the biofilms formed on the glass were gently washed twice with sterile BM base medium, and 400 μl of BM containing 20 mM Glc, GlcN, or GlcNAc was added back to the wells. After another 24 h of incubation, the spent medium was removed and the biofilms were washed three times with sterile PBS.

The washed biofilms were treated with a LIVE/DEAD BacLight bacterial viability reagent (Thermo Fisher Scientific, Waltham, MA), such that live bacteria produced green fluorescence (SYTO 9) and dead cells emitted red fluorescence (propidium iodide, PI). The architecture of the biofilms was examined using a confocal laser-scanning microscope (CLSM) (Leica, Wetzlar, Germany) equipped with a 60X oil immersion objective lens. VoxCell software (VisiTech International, Sunderland, United Kingdom) was used to acquire and process biofilm images. For SYTO9, the excitation wavelength was set at 488 nm and emission wavelength at 525 ± 25 nm; for PI, excitation was set at 642 nm and emission observed at 695 ± 53 nm. Biofilm stacks were also rendered in three dimensions using Imaris software (Bitplane, Belfast, United Kingdom).

To quantify bacterial viability in a dual-species biofilm, the biofilm was scraped off the coverslips, resuspended in sterile PBS and dispersed by sonication (FB120; Fisher Scientific) (30 sec × 2, 65% power). After serial dilution, bacterial suspensions were spread onto BHI agar plates with or without 1 mg ml^−1^ kanamycin (Sigma) to enumerate *S. mutans* and total bacteria, respectively.

### Measurement of H_2_O_2_ and Arginine deiminase (AD) activities

Levels of H_2_O_2_ in liquid TY cultures were measured by following an established protocol detailed elsewhere (14), and AD activity in bacterial cells was determined as previously described (58).

### Plate-based competition assays

Overnight bacterial cultures in BHI were washed with sterile PBS, resuspended in fresh TY medium containing 20 mM of the specified carbohydrate, with the OD_600_ being adjusted to 0.5. The cell suspensions (6 μl) of *S. mutans* UA159 and each commensal were spotted in close proximity on TY agar plates formulated with 20 mM Glc, GlcN or GlcNAc. Plates were incubated in aerobic or anaerobic conditions for 24 h at 37°C before being photographed.

### Bacteriocin production assays

Secretion of bacteriocins by *S. mutans* on agar plates was assessed according to a protocol detailed elsewhere (14), with minor modifications. Briefly, overnight cultures of *S. mutans* UA159 and commensal strains grown in BHI were washed with sterile PBS, and resuspended to an OD_600_ of 0.5 using fresh TY base medium supplemented with the desired carbohydrate. *S. mutans* UA159 alone or together with one commensal strain (at a ratio of 1:1) was inoculated by stabbing onto TY agar plates containing 20 mM Glc, GlcN or GlcNAc. After 24 h of incubation at 37°C, each plate was overlaid with 8 ml of warm soft agar (0.75% TY agar, prepared with the same sugar as was in the plate) containing 10^7^ cells of *S. sanguinis* SK150 as an indicator strain. Plates were incubated for an additional 24 h before examination for zones of inhibition of growth of *S. sanguinis*.

### Microcosm *ex vivo* biofilm model

Cell-containing saliva (CCS) samples (59) were collected and prepared from four healthy adult volunteers, who were non-smokers and had not taken antibiotics for at least 3 months (IRB201500497 at University of Florida). Equal volumes of untreated whole saliva from each volunteer were then pooled, glycerol was added to a final concentration of 25% and the CCS was stored in aliquots at −80°C. The pooled CCS was used as an inoculum for biofilm development by thawing of an aliquot and diluting 1:50 into biofilm medium (BM). When needed, exponential-phase cultures of *S. mutans* UA159-Km were prepared in BHI broth and inoculated at a ratio of 1:1000. For the biofilm experiments, BM medium was formulated with 2 mM sucrose and 18 mM glucose (BMGS), GlcN (BMGlcNS), or GlcNAc (BMGlcNAcS). Biofilms were allowed to form in Ibidi glass plates (Ibidi USA Inc., Madison, WI) at 37°C in an aerobic atmosphere supplemented with 5% CO_2._

Four different biofilm treatments were employed as detailed in Fig. 7A. Mixed cultures of *S. mutans* UA159-Km and CCS (group *a*), UA159-Km alone (group *b*), or CCS alone (group *c*) were inoculated into BMGS and incubated for 24 h. After washing with sterile, sugar-free BM medium, cultures were resupplied with BMGS, BMGlcNS or BMGlcNAcS, followed by 24 h of incubation. For group *d*, mixed cultures of UA159-Km and CCS were inoculated into BMGS, BMGlcNS or BMGlcNAcS. After 24 h of incubation, the supernates were replaced with the same, fresh media and the biofilms were incubated for another 24 h before harvesting.

To study the nature of the antagonistic effectors in these biofilms, CCS was used as the inoculum for biofilm development in BMGS, BMGlcNS or BMGlcNAcS for 24 h. At the same time, a panel of 24-h UA159-Km biofilms were established in Ibidi plates using BMGS, as described above. Filter-sterilized supernates (FSS, filter diameter 0.22 μm) were each prepared from the spent media of CCS biofilms and divided into three equal parts: 1) FSS supplemented with 1% catalase; 2) FSS passed through an Amicon^®^ filter with a molecular weight cut-off of 10 kDa; and 3) untreated FSS. UA159-Km biofilms were washed twice with PBS, then the various FSS samples were added and the biofilms were incubated for 24 h before harvesting.

To quantify the persistence of UA159-Km after these treatments, the biofilm cultures were washed twice with PBS, mechanically removed by scraping and resuspended in 400 μl of PBS, followed by sonication (30 sec × 2) to disperse the cells. The bacterial suspensions were then serially diluted, and plated on BHI agar plates containing kanamycin. All the plates were incubated for 48 h before CFU were counted.

### Statistical analysis

One-way ANOVA was performed to evaluate the significance of the comparisons from various experiments. The level of significance was determined at *P* <0.05.

## ACKNOWLEDGMENTS

This work was supported by DE12236 and DE25832 from the National Institute of Dental and Craniofacial Research. We thank Tanaz Farivar for technical support. LC was supported in part by a grant from China Scholarship Council (CSC).

